# Genotype-first approach revealed unrecognized breed-specific genetic diseases in dogs

**DOI:** 10.1101/2025.06.16.659006

**Authors:** Keijiro Mizukami, Ryoko Yamada, Hiroto Toyoda, Tomomi Aoi, Mikiko Endo, Junna Kawasaki, Ko Nakashima, Takaho Endo, Takashi Watanabe, Namiko Ikeda, Muneki Honnami, Naoki Fujita, Takayuki Nakagawa, Ryohei Nishimura, Jumpei Uchiyama, Masaharu Hisasue, Hajime Tsujimoto, Masahiro Sakaguchi, Hirotaka Tomiyasu, Yukihide Momozawa

## Abstract

Numerous dogs are affected by genetic diseases because of their unique breeding situation. The identification of pathogenic variants in dogs not only has veterinary benefits but also leads to the development of new human disease models. However, fewer pathogenic variants have been identified in dogs than in humans, and standard genetic approaches, such as large-scale case/control studies and family analyses, often fail to identify novel causative variants in dogs. To address this, we applied a genotype-first approach by developing a multiplex polymerase chain reaction (PCR)-based targeted sequencing method for dogs, analyzing 203 genes—most of which are related to human disease—from 6,333 dogs collected from a veterinary hospital, and characterizing pathogenic variant carriers. In total, 120 pathogenic variants in 83 genes were identified. Frequent pathogenic variants were enriched in specific dog breeds. Dogs homozygous for a pathogenic *CHEK2* variant, frequently identified in French Bulldogs (5.74%, homozygous), showed a five-fold increased risk of cancer (*P* = 3.45 x 10^-2^). A pathogenic *NOD2* variant was detected in half of the Italian Greyhounds. However, their medical records did not document the typical clinical symptoms observed in humans with *NOD2* deficiency. Some pathogenic variants were present in three or fewer dogs, and 11.0% of these dogs exhibited symptoms potentially associated with gene defects. These rare variants could become common within a short period due to the canine breeding system to induce breed-specific genetic diseases. These findings suggest that the genotype-first approach successfully identified unrecognized breed-specific genetic diseases. Further characterization of dogs with pathogenic variants enable to define clinical diagnosis, providing an important spontaneous model for human genetic diseases.

## Introduction

The discovery of genetic diseases in domestic dogs has led to the development of novel spontaneous models because many genetic diseases in dogs have a similar clinical presentation to homologous human diseases and are enriched in specific breeds owing to strong selection during breed development (1). Canine genetic diseases that have already been identified serve as important models for studying rare human diseases. For example, the efficiency of antisense oligonucleotides for Duchenne muscular dystrophy was confirmed in Golden Retrievers with pathogenic variants in *DMD* (2), which finally led to the development of drugs (viltolarsen) for the disease in humans (3). Their relatively long lifespan compared to general experimental animals (e.g., mice) also makes them useful as models for studies requiring long-term observation; for example, studies have been conducted in hemophiliac dogs to elucidate the long-term efficacy of adeno-associated viral vector gene therapies in patients with hemophilia (4) and the mechanisms of vector persistence in the liver (5). In addition, because dogs are companion animals, it is important to free them from suffering because of diseases. Identification of the pathogenic variant of a genetic disease allows genetic testing to be established, thus reducing the number of dogs with the disease based on breeding management (6). Therefore, the identification of causative genes and variants of genetic diseases in dogs is important from both medical and veterinary perspectives.

Genes with variants associated with genetic diseases have been identified in dogs. Currently, the animal genetic disease database OMIA contains information on 323 phenotypes of canine genetic diseases with 502 deleterious variants, most of which are breed-specific (accessed December 19^th^, 2024) (7). However, the number of genetic diseases discovered in dogs is approximately 10% of that in humans, according to the human genetic disease database OMIM (accessed on December 19^th^, 2024), and most diseases for which causative genes and variants have been identified are limited to those caused by pathogenic alleles with high penetrance (8). Most variants associated with genetic diseases have been identified by sequencing or genotyping using large-scale case-control studies or family analyses of probands in humans (9). Although standard genetic approach (e.g., genome-wide association studies) generally require very large sample sizes to identify reproducible significant associations (10), it takes a long time to collect sufficient case and control samples from the same breeds diagnosed under homogenized diagnostic criteria. In addition, epidemiological data are useful for obtaining preliminary information for studying inherited diseases (e.g., heritability and inherited patterns), which are often unavailable in case-control studies of canine diseases. Furthermore, the inaccessibility of the family history of pet dogs isolated from blood relatives makes it difficult to estimate whether a genetic background exists for these symptoms. Therefore, an approach different from standard genetic one is required to address this issue.

To address this difficulty, we applied genotype-first analyses, detecting pathogenic variants in disease-associated genes in all patients used for data collection without selection by clinical presentation, and analyzed the association between the identified variants and clinical information to detect potential genetic diseases (11). This approach eliminates the phase of preselecting conditions suspected to be hereditary, allows sample collection without requiring a detailed diagnosis at the time of sampling, and does not require the collection of pedigree information. In this study, DNA samples were obtained from the blood test residuals of over 6,300 canine cases presenting at a veterinary hospital, and multiplex PCR-based targeted sequencing, which is the most high-throughput method for analyzing specific genes, was developed for dogs. Using these samples and methods, we sequenced all coding regions and 2-bp flanking intronic sequences of 203 genes, mostly related to human diseases, identified pathogenic variants in these genes, and analyzed the clinical information of dogs carrying these variants.

## Results

### Study design and sample collection

The experimental and analytical flow of this study is shown in Fig. 1A. In the present study, we analyzed 203 genes. Of these, 61 genes were from the ACMG Recommendations for Reporting of Secondary Findings in Clinical Exome and Genome Sequencing (12) and 121 genes associated with various human diseases in previous reports were included to evaluate the possibility of using dogs as a model for human hereditary diseases (Fig. 1B, SI Appendix, Table S1). The remaining 21 genes were associated with morphology or blood test values in dogs; 12 of them were also related to human diseases, and the others were included to analyze potential disease associations. The dogs analyzed in this study were not limited to patients from a specific clinical department; therefore, genes associated with various diseases were included to cover a wide range of conditions.

**Figure 1.**
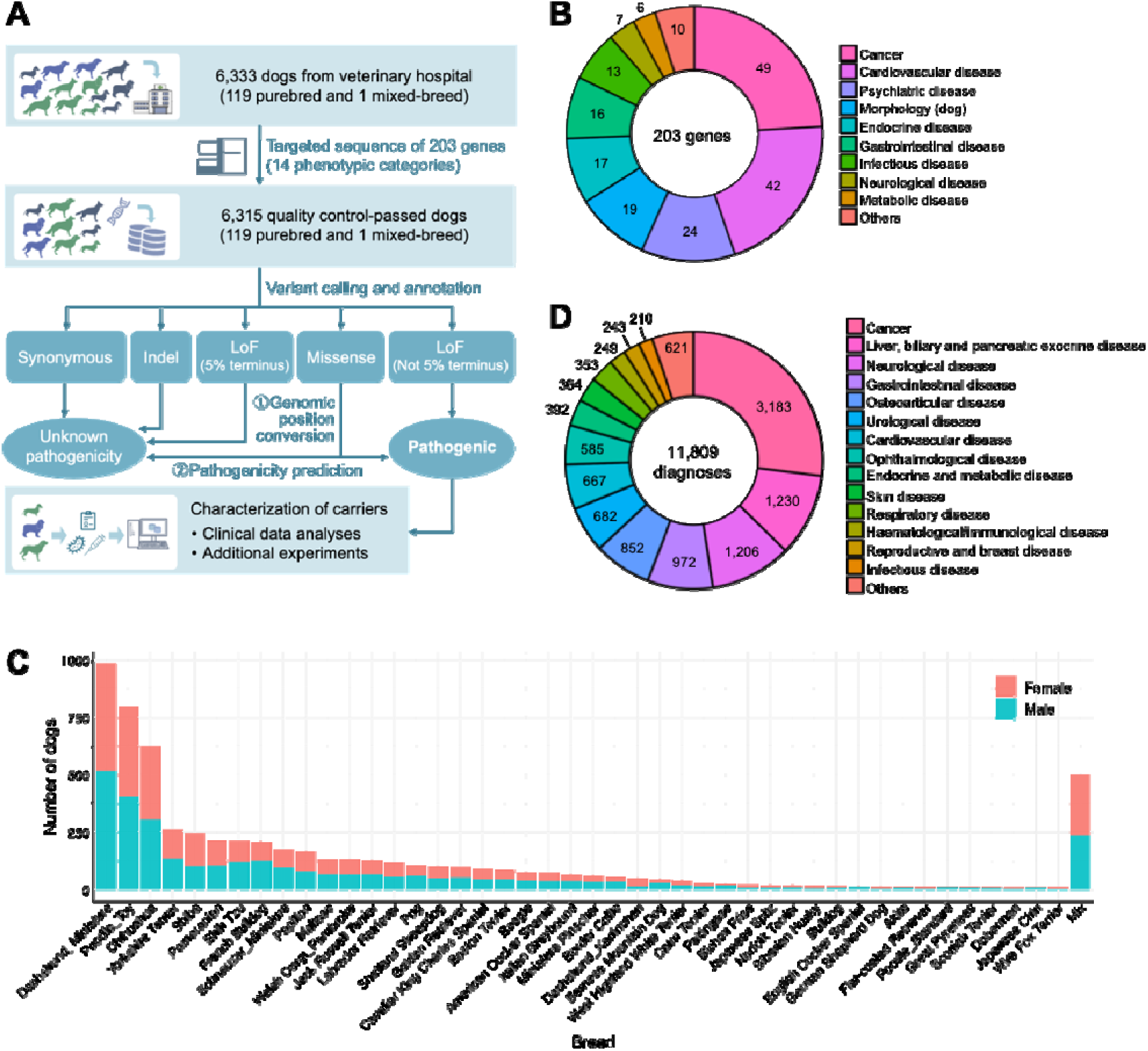
Overview of this study, targeted gene, and samples. (A) Flow of this study. (B) The composition of the diseases category of targeted genes. The phenotypic category of “others” in targeted genes included musculoskeletal disorder (n = 4), blood test parameter in dogs (2), dental (2), respiratory (1), and urological (1) disease. (C) Breed distribution in the study population (only breeds with ≥ 10 dogs). (D) The composition of the diagnoses of samples analyzed in this study. The diagnoses of “others” included oral disease (n = 189), drug-induced condition (113), allergies (87), undiagnosed/unclassifiable (80), trauma/accidental ingestion (64), behavioral disease (62), muscular disease (19), and conditions associated with surgery (7).

A total of 6,333 dogs from 119 breeds (3,183 males and 3,150 females) were analyzed in this study (Fig. 1C; SI Appendix, Table S2). DNA was extracted from residual EDTA blood of patients who presented to the Veterinary Medical Center of the University of Tokyo and underwent blood testing. The most abundant breeds were Toy Poodles, Miniature Dachshunds, and Chihuahuas, which is consistent with the prevalent dog breeds in Japan. Of the 6,333 dogs, 5,823 had at least one clinical diagnosis. Because most dogs had multiple diagnoses, the total number of diagnoses was 11,809. The most frequent diagnosis was cancer (3,183 cases), followed by liver, biliary, and pancreatic diseases (1,230 cases) and neurological diseases (1,206 cases) (Fig. 1D).

### Development of multiplex PCR–based targeted sequencing

Multiplex PCR–based targeted sequencing was successfully established for dogs. For each gene, the average proportions of regions with 20x coverage within the target region were 98.6% (standard deviation [SD], 2.78). The average proportion of regions with 20x coverage across the entire target region for each sample was 98.4% (SD, 1.76). Following sequencing quality control procedures, we analyzed 6,315 dogs (3,138 males and 3,150 females) from 119 breeds that had at least 95% of the target region with 20x coverage. We identified 3,918 variants. The number of variants per gene showed a significant correlation (Spearman’s ρ = 0.772, *P* = 1.32 × 10^-40^) with the total length of the targeted gene region (SI Appendix, Fig. S1). The number of variants detected per breed was strongly correlated with the number of dogs in each breed (Spearman’s ρ = 0.938, *P* = 1.51 × 10^-55^) (Fig. 2A). There was no clear bias toward a particular clade (13). The number of non-synonymous and synonymous alleles was small in Boxers because this breed was used for the reference genome (CanFam3.1) in this study. These results suggest the reasonable performance of variant calling in this sequencing approach. Among the 3,918 variants: 90 loss of function (LoF) variants, containing 10 LoF variants within 5% terminus of the protein, 1,733 non-synonymous variants, and 2,095 synonymous variants. The average number of LoF, non-synonymous and synonymous alleles detected in a dog was 6.98 x 10^-2^, 121, and 353, respectively (SI Appendix, Fig. S2).

**Figure 2.**
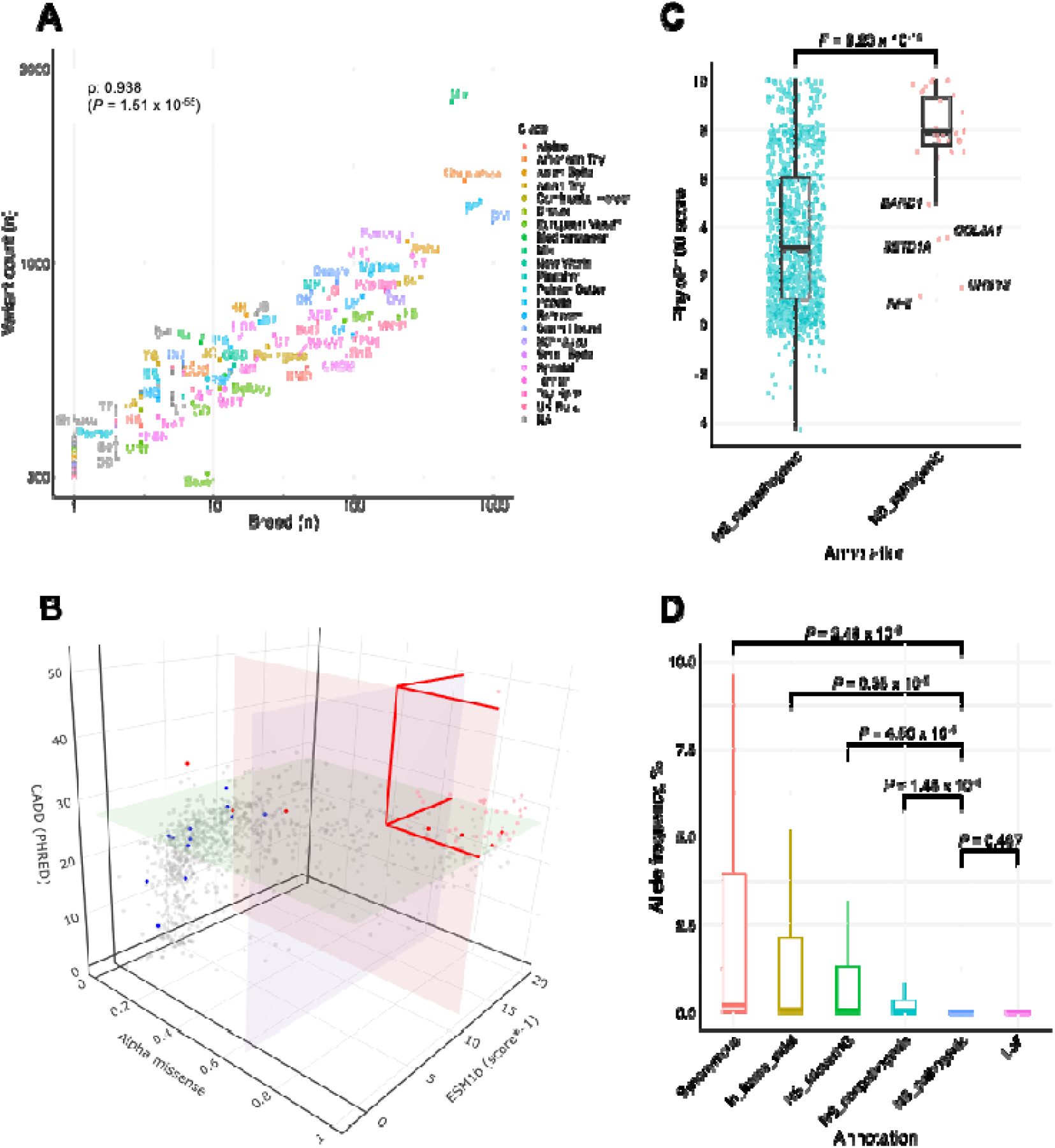
Characteristics of the variants detected in this study. (A) The correlation of the number of variants detected in each breed and the number of breeds. The upper and lower breed outliers were mixed dogs and Boxer (the breed of the reference genome), respectively. (B) 3D-plot shows the distribution of three *in silico* programs’ score of missense variants. The blue-, pink-, and green-colored panels represents the thresholds calculated in this study for AlpahMissense, ESM1b, and CADD scores, respectively. The pink-colored plots surrounded by red lines are variants annotated as pathogenic in this study. The red- and blue-colored plots are variants annotated as pathogenic/likely-pathogenic and benign/likely-benign in ClinVar. (C) Comparison of conservation between pathogenic missense variants (NS_pathogenic) and non-pathogenic missense variants the genomic positions of which were successfully lifted to the human genome (NS_nonpathogenic). NS_pathogenic were more conserved than NS_nonpathogenic. (D) Comparison of allele frequencies among each annotation. The allele frequencies of NS_pathogenic are shown with those of loss of function variants (LoF), synonymous variants, in-frame indel variants (In_frame_indel), missense variants in the genomic position of which could not be lifted to the human genome (NS_liftoverNG), and NS_nonpathogenic. Abbreviations of breed name are described in Table S2 in SI Appendix.

To confirm that this method detected known disease-associated variants in dogs, we examined the variants registered in OMIA. Of the 19 single nucleotide or short indel variants in the 203 genes registered in OMIA (last updated on December 19^th^, 2024), six variants were detected in this study: p.Gln187fs in *POMC* related to obesity (14), p.Cys188Phe in *SLC2A9* related to urolithiasis (15), p.Arg1384Gln in *ATP7B* related to Wilson disease (16), p.Phe665fs in *ENAM* related to amelogenesis imperfecta (17), p.GluLys154* in *APC* related to familial adenomatous polyposis (18), and p.Lys165del in *CARD9* related to susceptibility to *Mycobacterium avium* complex (19) (Table S4). All dogs (6/6) with variants in *APC* showed gastric and/or colorectal cancers, which are phenotypes observed in humans and dogs with familial adenomatous polyposis caused by the gene. Although body condition information was unavailable in this study, dogs with the variant in *POMC* were more frequently diagnosed with cancer (*P* = 8.96 x 10^-4^, Cochran-Armitage test), which may reflect conditions secondary to obesity (20). Only heterozygous dogs were found for variants in *ENAM* and *CARD9*. Since both genes followed an autosomal recessive inheritance pattern, the dogs did not show symptoms related to the variants. For the 13 missing variant regions, 98.5% of the samples were analyzed with adequate coverage (i.e. 20 and more sequencing reads), but no registered variants were identified, suggesting that they are monomorphic in our dogs. These results suggest that this genome-first approach has a high potential to identify pathological variants.

### Pathogenicity prediction of missense variants

Although LoF variants could be considered pathogenic if they exist in genes whose defects lead to disease (21), it is challenging to determine the pathogenicity of non-synonymous variants. In this study, we predicted the deleteriousness of the detected missense variants using stateloflthelart *in silico* programs, AlphaMissense (22), ESM1b (23), and CADD v1.7 (24). Of the 1,733 non-synonymous variants detected in this study, 1,644 were missense variants, and 1,107 were variants whose genomic positions were successfully lifted from CanFam3.1 to hg38, confirming the consistency of amino acids before and after substitutions between dogs and humans. The variants were scored using the three *in silico* programs. Forty missense variants in 36 genes had more deleterious scores than the predefined thresholds for all programs and were considered pathogenic variants in this study (Fig. 2B).

To confirm the relevance of the characteristics of the pathogenic missense variants, we compared the conservation and allele frequencies of the variants with those of variants with different annotations. Based on the hypothesis that the genomic positions of pathogenic missense variants are more conserved, we compared the PhyloP scores between pathogenic missense variants and missense variants whose positions were lifted to hg38 and deleteriousness scores did not exceed the threshold in at least one of the *in silico* programs (NS-nonpathogenic). The PhyloP scores for pathogenic missense variants were higher than those for NS_nonpathogenic variants (*P* = 9.23 x 10^-14^, Wilcoxon rank-sum test) (Fig. 2C). When comparing the allele frequencies of pathogenic missense variants with other annotations, there was no difference between pathogenic missense and LoF variants (*P* = 0.467, Wilcoxon rank-sum test) while the allele frequencies of pathogenic missense variants was significantly lower than those of variants with other annotations (*P* < 0.01 = 0.05/5) (Fig. 2D). These results showed that pathogenic missense variants tend to be located in evolutionarily constrained regions and, like LoF variants, have a negative effect on survival.

### Characteristics of pathogenic variants

Except for the LoF variants at the 5% terminus of the protein, 80 LoF variants were detected in 57 genes. In addition to the 40 pathogenic missense variants detected in 34 genes, 120 variants in 83 genes were defined as “pathogenic variants” (SI Appendix, Fig. S3, Table S3). The highest cumulative allele frequency of pathogenic variants at the gene level was observed for *POMC*, followed by *BARD1* and *CHEK2* (Fig. 3A). A comparison of allele frequencies in dogs with Shet scores, which quantify fitness loss because of heterozygous LoF variation (25), showed a distribution similar to that of the 203 genes analyzed in humans (Fig. 3B; SI Appendix, Fig. S4), indicating that genes that are less prone to harbor LoF variants are shared between the two species. The correlation of the cumulative allele frequencies of LoF variants between dogs and humans (obtained from gnomAD (26)) was weak (Spearman’s ρ = 0.348, *P* = 8.93 × 10^-7^) and both higher- and lower-frequency genes were also present (Fig. 3C). In particular, the allele frequencies of LoF variants in *NOD2*, *POMC*, *APOB*, *CHEK2*, *ACP4*, *LCORL,* and *ENAM* were 4.98 (*ACP4*) to 36.0 (*LCORL*) times more frequent in dogs than in humans.

**Figure 3.**
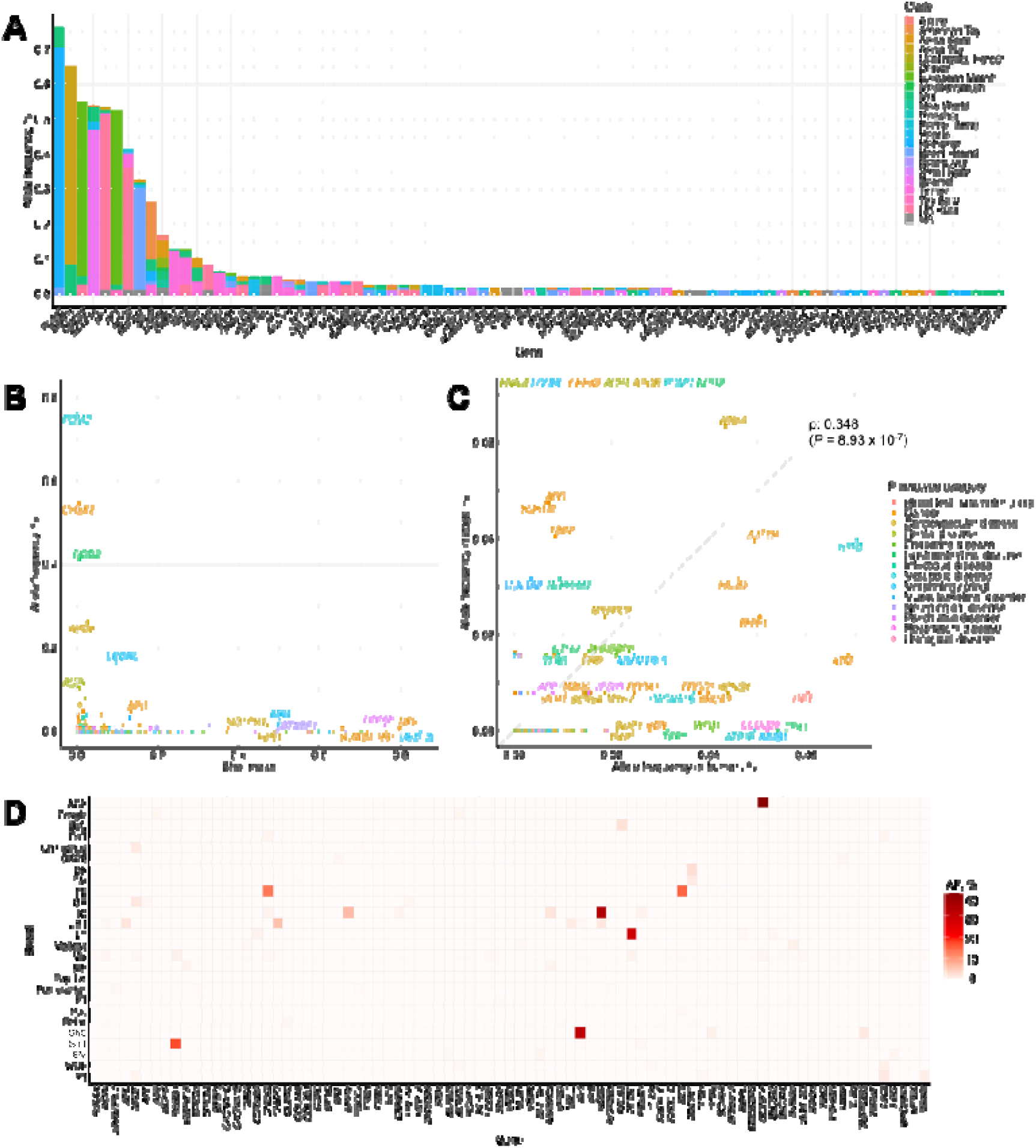
Characteristics of genes with pathogenic variants. (A) The cumulative alternative allele frequencies of pathogenic variants at the gene level. The colors of each bar represent the clades to which the breeds of the carrier dogs belong to (13). (B) Distribution of the alternative allele frequencies in dogs and the mean Shet scores, which quantifies fitness loss because of heterozygous LoF variation in humans (25), of each gene. The distribution showed similar pattern of that in humans. (C) Comparison of the allele frequencies of LoF variants at the gene level between humans and dogs. Weak correlation of the alternative allele frequencies of each gene between humans and dogs were observed. (D) The matrix of the allele frequencies of pathogenic variants at gene level by breed with over 50 dogs. AF, allele frequency.

High-frequency pathogenic variants were enriched in specific breeds (Fig. 3D; SI Appendix, Fig. S5). In breeds with over 50 dogs, a >10% allele frequency within breeds was observed for seven pathogenic variants. Of these variants, four were located in hereditary cancer-related genes: *NF1* (p.Ala1451Pro enriched in Shetland Sheepdog; allele frequency [AF] 31.2%), *BARD1* (p.Leu289Phe in Shih Tzu; 17.1%), *RET* (p.Val256Met in French Bulldog; 15.1%), and *CHEK2* (p.Leu184fs in French Bulldog; 13.4%). The pathogenic missense variant of *SETD1A* (p.Arg106Trp), associated with schizophrenia, was enriched in American Cocker Spaniels (40.4%). The LoF variant in *NOD2* (p.Val928fs), a gene associated with Crohn’s disease, was enriched in Italian Greyhounds (33.3%). The LoF variant in *POMC* (p.Gln187fs) was associated with obesity in Labrador Retrievers, but the allele frequency in the breed used in this study (28.1%) was higher than that reported in a previous study (12%) (14).

### Analyses of LoF variants in *CHEK2* and *NOD2*

We focused on the two most prevalent and newly identified LoF variants. The clinical characteristics of LoF variants of *CHEK2* (p.Leu184fs) and *NOD2* (p.Val928fs), which are frequently detected in French Bulldogs and Italian Greyhounds, respectively, were investigated. The LoF variant of *CHEK2* was located in the first half of the gene and was predicted to completely disrupt the protein kinase domain (Fig. 4A). *CHEK2* is a hereditary cancer-related gene known to be associated with a two-to three-fold increase in the risk of multiple cancers, including female breast and prostate tumors (27); therefore, the association between the presence of cancer and the genotypes of the variant was investigated based on clinical diagnosis. Dogs homozygous for the variant (5.74% of French Bulldogs) were more likely to have cancer than heterozygous dogs and those homozygous for the wild-type allele (odds ratio 5.02, *P* = 3.45 x 10^-2^, Fisher’s exact test under recessive model) (Fig. 4B). When investigating the cancer types, dogs homozygous for the variant showed a nominal association with hepatic tumors (*P* = 4.00 x 10^-2^, Fisher’s exact test under recessive model), and the frequency of cases with brain and mast cell tumors showed an additive increasing trend with respect to the number of LoF variant alleles (SI Appendix, Fig. S6).

**Figure 4.**
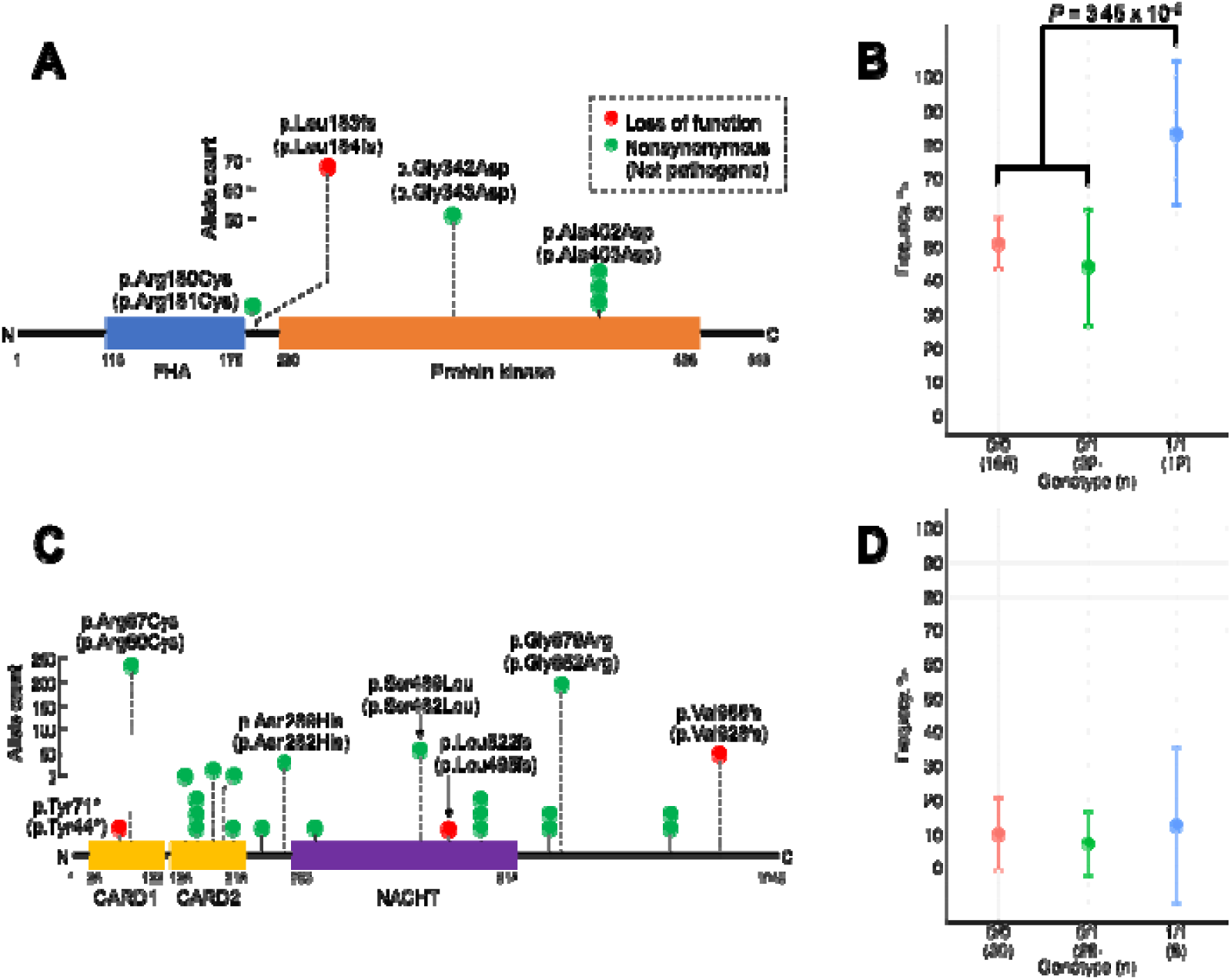
Characteristics of the LoF variants in *CHEK2* and *NOD2*. (A) Location and number of variants in *CHEK2* detected in this study are shown. Because the domains information was derived from human CHKE2 (UniPro ID: O96017) obtained from UniProt, the positions are based on human protein. The positions of variants in dogs are indicated in parentheses. (B) Comparison of frequencies of dogs with cancers among the genotypes of the LoF variant in *CHEK2* (p.Leu184fs). Plots and bars represent averages and standard errors of these frequencies. Dogs homozygous for the variants were more likely to be diagnosed with cancer than other genotypes. (C) Location and number of variants in *NOD2* detected in this study are shown. Because the information for the domains was derived from human NOD2 (Q9HC29) obtained from UniProt, the positions are based on human protein. The positions of variants in dogs are indicated in parentheses. (D) Comparison of frequencies of dogs with gastrointestinal symptoms among the genotypes of the LoF variant in *NOD2* (p.Val928fs). The frequencies were comparable among genotypes of the variant. FHA, forkhead-associated; CARD, caspase recruitment domains.

The LoF variant in *NOD2* (p.Val928fs), frequently observed in Italian Greyhounds, was present in the latter part of the protein and predicted to create a premature stop codon after 14 amino acids (Fig. 4C). Frameshift variants at the 1,005th and 1,007th codons of the 1,040 amino acids of the wild-type protein in humans have been reported to be associated with Crohn’s disease (28, 29). Therefore, we examined the presence of gastrointestinal symptoms, including diarrhea, in each genotype of the variant, but no association was observed between the presence of symptoms and genotypes (Fig. 4D). Although *NOD2* has also been reported to be associated with resistance to bacterial infections (30), none of the dogs had a diagnosis of bacterial infection in their medical history. We also checked the number of bacteria-derived sequence reads in the peripheral blood for each genotype from 47 whole-genome sequencing datasets to explore potential bacterial infections. Bacteria, including Pseudomonadota, Bacillota, Actinomycetota, Bacteroidota, and Bacteroidota/Chlorobiota, were detected. However, no genotype-dependent changes were observed in the presence of specific bacteria (SI Appendix, Fig. S7A), the total read number of bacteria (SI Appendix, Fig. S7B), or susceptibility to specific bacteria (SI Appendix, Fig. S7C), without contradicting the clinical information. No virus infection was observed. These data indicate that the variants did not affect the bacterial and virus composition of the blood.

### Pathogenic variant carriers in other genes

In addition to the pathogenic variants enriched in specific breeds, several variants detected in this study were detected in only a small number of cases. Ninety-four pathogenic variants from 65 genes were present in three or fewer dogs, 51 of which were variants in genes that caused the disease in a dominant pattern. Of these 94 variants, 73 variants in 54 genes were identified in one dog, and 21 variants in 17 genes were identified in two or three dogs. Of the dogs with these variants, 13 (11.0%) were affected by diseases that were assumed to be related to genetic defects. For example, a heterozygous LoF variant in *MSH6* (p.Pro1010fs) has been identified in a Shiba affected by intestinal cancer. Two LoF variants in *BRCA2* (p.Ser750fs and p.His744fs), which are associated with breast cancer, were also identified in a female Chihuahua. A pathogenic missense variant (p.Arg846His) of *GRIN2A*, an epilepsy-related gene, was detected in a mixed female dog diagnosed with epilepsy.

## Discussion

In this study, we developed a multiplex PCR–based targeted sequencing method for dogs and performed a genotype-first approach by analyzing 203 mainly human disease-causing genes in 6,315 dogs collected from a veterinary hospital. Of the 3,918 variants identified, 120 were pathogenic variants as defined in this study. Frequent pathogenic variants have been identified in specific breeds. Dogs homozygous for an LoF variant of *CHEK2* have an increased risk of cancer, while dogs with an LoF variant in *NOD2* did not show the typical clinical symptoms observed in humans with *NOD2* deficiency in their medical records. These results indicate that unrecognized breed-specific diseases may remain latent in dogs.

Pathogenic variants are frequently observed in certain breeds, leading to unrecognized, breed-specific diseases. Cancers related to *CHEK2* have not been reported in non-human, non-experimental animals so far. An LoF variant of *CHEK2* has been newly identified in brachycephalic breeds, including French Bulldogs, Bulldogs, and Boston Terriers, in this study. An increased risk of developing a broad range of cancers, including breast, colorectal, and prostate cancers, has been observed in human patients with *CHEK2* deficiency. However, further data are required to confirm clinically actionable associations with other types of cancer (27). In contrast, other cancers, including rare cancers in humans (e.g., mast cell tumors), showed a trend toward an association with the genotypes of the LoF variant of *CHEK2* in dogs. This may be because of differences in the general incidence rates of each cancer type between humans and dogs (31). Because dogs homozygous for the variant were five times more likely to develop cancer than dogs heterozygous or homozygous for the wild-type allele, it is important to gradually reduce the variant in the French Bulldog population through breeding management. In addition, because homozygotes or compound heterozygotes of LoF variants are rare in humans, homozygous dogs could be a crucial model for cancers caused by multiple LoF variants.

A few or singleton cases with pathogenic variants in disease-related genes were observed in this study. Dogs harboring LoF variants of *BRCA2* and *MSH6,* which are hereditary cancer genes, developed breast and intestinal cancers, respectively. As the carrier frequencies of LoF variants in *BRCA2* and *MSH6* in human non-cancer populations are 0.17% and 0.09%, respectively (32), the carrier frequencies (0.0158%) in the dog population in this study, which contained a large number of cancer cases, were lower than those in humans. This may be because of natural and/or artificial selection in canine breeding system. On the other hand, some LoF variants detected in a small number of dogs were found in genes with high Shet scores, where carrier frequencies are lower in humans (25). For instance, LoF variants in *NF1* (Shet 0.797, AF in humans 8.76 × 10l³%) and *TRRAP* (0.761, 1.46 × 10l³%), associated with neurofibromatosis type 1 and developmental delay, respectively, were identified in singleton dog cases, highlighting their potential value in providing novel insights into rare human diseases despite their rarity in dogs. In addition, it should be noted that these rare variants are likely to become common within a short period in the future because of popular sire and founder effects in specific regions (33) to induce breed-specific genetic diseases.

Of the 120 pathogenic variants, three were known disease-associated variants, nine were in genes in which other variants had already been reported in dogs, and 108 were identified in genes in which no disease-associated variants in dogs had been registered in the OMIA. Although some dogs harboring pathogenic variants exhibited clinical symptoms that may have resulted from alterations in gene function, others did not exhibit such symptoms. Although not all genes analyzed in this study showed complete penetrance of the relevant disease, there may be several reasons for the absence of related clinical symptoms in the carrier dogs. Some carrier dogs may not have reached the age of disease onset associated with these genes. For example, an LoF variant in *BRCA1* was detected in a female Pomeranian without a diagnosis of any cancer, which was one year old at the time of the last recorded clinical data used in this study. Since carriers of pathogenic variants develop cancer after approximately 45 years of age in humans (34), carrier dogs could develop cancer in the future. Other possible reasons are that phenotypes related to these genes might be undetectable in veterinary practice or might not exist in dogs because of physiological differences between species and the absence of organs because of ovariohysterectomy and subsequent hormonal imbalance in dogs. Therefore, continuous clinical observation and detailed biological analyses of carrier dogs are required using samples from these dogs.

In conclusion, this study identified 120 pathogenic variants in canine genes homologous to human disease-related genes using a genotype-first approach. Most of these variants were newly discovered, highlighting the potential presence of unrecognized genetic diseases in dogs. These findings expand the scope of genomic medicine in the veterinary field and demonstrate the broader applicability of the genotype-first approach to non-human, non-experimental animals.

## Materials and Methods

### Samples collection

DNA from 6,333 dogs of 119 breeds (3,183 males and 3,150 females) was extracted from the residual EDTA blood of patients who presented to the Veterinary Medical Center of the University of Tokyo and underwent blood tests. Clinical information of the dogs was collected from hospital medical records. Informed consent was obtained from the owners of all the participants. Ethical approval was obtained from the RIKEN Center for Integrative Medical Sciences (AEY2024-028).

### Development of multiplex PCR–based targeted sequencing and variant calling

Multiplex PCR-based targeted sequencing has been developed for dog genomes. Primers for the coding regions and 2-bp flanking intronic sequences were designed based on CanFam3.1 using Primer 3 (ver. 2.3.6) (35) to avoid known variants in the Ensembl SNPs database (accessed on February 23^rd^, 2019) and the 170lK Illumina HD canine SNP array (36). Exon regions were determined based on the annotation of all isoforms in RefSeq, although only the annotation of curated isoforms (i.e., transcript ID starting with NM) was used, if available. The procedures for multiplex PCR–based targeted sequencing and variant calling for the dog genome were modified from those developed for the human genome in our laboratory (37). Detailed information is provided in the SI Appendix.

### Variant annotation and determination of pathogenic variants

In this study, LoF variants that were not located in the 5% terminal region of the protein and missense variants predicted to be deleterious using *in silico* programs were considered pathogenic variants. Each variant was annotated using SnpEff (ver. 4.3) (38) with transcripts used in primer design. Although determining whether missense variants are pathogenic is challenging, high-precision programs have been developed (22–24). In this study, we focused on missense variants in the genomic position that were successfully lifted from dogs to humans and adapted only consistent prediction results among three state-of-the-art programs to increase specificity: AlphaMissense (22), ESM1b (23), and CADD v1.7 (24). Detailed information on variant annotation is provided in the SI Appendix. To determine the thresholds of deleteriousness scores for each program, ClinVar’s pathogenicity annotations were predicted for variants in 203 targeted genes registered in the database using the score of each program. All programs showed high prediction accuracies (ROC-AUC score: AlphaMissense, 0.972; ESM1b, 0.940; CADD v1.7, 0.919) (SI Appendix, Fig. S8). The scores with maximum true positive rate among false positive rates of 5% or less were 0.6595 for AlphaMissense, -11.121 for ESM1b, and 27.6 for CADD v1.7 (PHRED); hence, these values were used as the thresholds of deleteriousness in this study. To confirm the genomic conservation of missense variants annotated as pathogenic, phyloP100way scores at the human genome position (hg38) lifted from the canine genome position (CanFam3.1) of the missense variants were obtained using ANNOVAR (v2020-06-07) (39).

### Further phenotypic analyses of LoF variant in *NOD2*

To characterize dogs with each genotype of the identified LoF variant in *NOD2*, we explored the potential phenotypes of *NOD2* deficiency. The number of bacteria-derived sequence reads in the blood were counted using unmapped sequence reads in whole-genome sequencing of DNA derived from the peripheral blood of 47 Italian Greyhounds (21 homozygotes and 20 heterozygotes of the wild-type allele and 6 homozygotes of the alternative allele). Unmapped sequence reads were separated from whole-genome sequencing data and aligned to the reference sequence (UU_Cfam_GSD_1.0) using the Burrows-Wheeler Aligner (ver. 0.7.9a) (40) and taxonomic diversity was assessed using STAT (41).

### Data availability

The multiplex PCR–based targeted sequencing data from 6,315 dogs and unmapped sequencing data from whole genome sequencing in 51 Italian Greyhounds are publicly available in the DDBJ (PRJDB20692).

## Acknowledgments

We sincerely appreciate the dog owners, veterinarians, and veterinary hospital staff for their invaluable support in providing the clinical samples for this study. We also extend our gratitude to the students and staff of the School of Veterinary Medicine at Azabu University for their assistance in DNA extraction. Additionally, we thank the Genomics and Transcriptomics Unit and the Advanced Multi-Omics Technology Division at the RIKEN Center for Integrative Medical Sciences for their support. This work was supported by JSPS KAKENHI Grant Number 22K18377 and JP23K05584 and by Ministry of Education, Culture, Sports, Science and Technology (MEXT) Supported Program for the Private University Research Branding Project, 2016–2020.

## SI Figure and Table legends

**Figure S1.** The correlation of the length of targeted genes and the number of the variants detected in this study.

**Figure S2.** The total number of variants detected per breed (bar plot with left y-axis) and the number of alleles per individual for each breed (line plot with right y-axis).

**Figure S3.** The alternative allele frequencies of pathogenic variants in bar plots. The colors of each bar represent the clades to which the breeds of the carrier dogs belong to Parker, et al. (13).

**Figure S4.** Distribution of the alternative allele frequencies in human (gnomAD) and the mean Shet scores of each gene.

**Figure S5.** The matrix of the allele frequencies of pathogenic variants by breed with over 50 dogs. Abbreviations of breed name are described in Table S2 in SI Appendix. AF, allele frequency.

**Figure S6.** Comparison of frequencies of dogs with each cancer among the genotypes of the LoF variant in *CHEK2* (p.Leu184fs). Plots and bars represent averages and standard errors of the frequencies. Hepatic cancer showed nominal association (*P* = 4.00 x 10^-2^) with the genotype of the variants at the recessive model.

**Figure S7.** The estimated number of bacteria in the peripheral blood for each genotype of the LoF variant in *NOD2* using unmapped sequence reads from 51 whole-genome sequencing data. No genotype-dependent alternation in the presence of certain bacteria (A), the total number of bacteria (B) or susceptibility to certain bacteria (C) were observed.

**Figure S8.** Receiver operating characteristic curves showing predictive performance of scores from three *in silico* programs with ClinVar annotation (pathogenic, likely_pathogenic, benign, and likely_benign) of variants in genes analyzed in this study.

**Table S1.** Information of targeted genes in this study. ACMG, American College of Medical Genetics. OMIM, Online Mendelian Inheritance in Man.

**Table S2.** Sample number, abbreviation, and clade of breeds analyzed in this study.

**Table S3.** Annotation of pathogenic variants detected in this study. Positions of HGVS.c and HGVS.p were based on the longest annotation among isoforms in RefSeq although only the annotation of the curated isoforms were used when available. LoF, loss of function variant. In_frame_indel, in-frame indel variant. NS_liftoverNG, missense variants of which genomic position could not be lifted to the human genome. NS_nonpathogenic, non-pathogenic missense variants of which genomic position was successfully lifted to the human genome. NS_pathogenic, missense variants annotated as pathogenic.

**Table S4.** OMIA-registered variants in targeted genes in this study. OMIA, Online Mendelian Inheritance in Animals. LoF, loss of function variant. In_frame_indel, in-frame indel variant. NS_liftoverNG, missense variants of which genomic position could not be lifted to the human genome. NS_nonpathogenic, non-pathogenic missense variants of which genomic position was successfully lifted to the human genome.

