## Supplemental appendix for "Genotype-first approach revealed unrecognized breed-specific genetic diseases in dogs"

### Supporting Methods

#### Development of multiplex polymerase chain reaction–based targeted sequencing and variant calling

The sizes of the PCR products of the multiplex PCR–based targeted sequencing were designed to be 180–295 bp to cover the amplicon with sequencing reads. We added CGCTCTTCCGATCTCTG to the 5′ end of the forward primers and CGCTCTTCCGATCTGAC to the 5′ end of the reverse primers to perform the second PCR (1). Multiplex PCR reactions were carried out in 10 µl containing 10 ng genomic DNA, 5 µl 2× Platinum Multiplex PCR Master Mix (Life Technologies), and 0.13 - 0.2 pmol from each primer. Cycling conditions were 95 °C for 2 min, 25 cycles of 95 °C for 30 s, 60 °C for 90 s, and 72 °C for 2 min, followed by an additional extension step at 72 °C for 10 min using the GeneAmp PCR System 9700 (Life Technologies). Primer sequences for the second PCR were 5′ -AATGA TACG GC GACCACCGAGA TCTAC ACxxxxxxxxACA CTC TT TC CCTA CACGAC GCTCTTC CGATCTCTG-3′ and 5′ -CAAGCAGAA G ACGGCATACGAG ATxxxxxxxxGTGAC TGGAGTTCAGACGTGTG CTC TTCC GATCTGAC-3′. We used 384 types of barcodes containing the first 8 bp from 10 bp of the barcode sequences reported by Forshew et al. (2). The PCR reactions were carried out in 10-µl reactions containing 2 µl of the first PCR product, 2.5 µl KOD One PCR Master Mix (TOYOBO), and 0.2 pmol from each primer. Cycling conditions were 94 °C for 5 min, 6 cycles of 98 °C for 10 s, 60 °C for 5 s, and 68 °C for 10 s. All second PCR products were pooled for one sequencing run. After each completed session, the library was purified using Agencourt AMPure XP (Beckman Coulter) to eliminate primer dimers, was applied to a bioanalyzer (Agilent Technologies) to check the size distribution and then quantified using the KAPA library quantification kit (KAPA) on an ABI Prism 7900HT sequence detection system (Life Technologies). We obtained 2 × 150-bp paired-end reads with dual 8-bp barcode sequences on a HiSeq 2500 instrument. A 'dark cycle' was applied to discard the first 3 bp of both reads originating from the end of the adapter sequences used for the second PCR.

Sequence reads were aligned to the dog reference sequence (CanFam3.1) using the Burrows-Wheeler Aligner (ver. 0.7.17) (3) and then applied to RealignerTargetCreator and IndelRealigner using GATK (ver. 3.7) (4) for each BAM file. For quality control, dogs were excluded from further analysis if the proportion of covered bases of ≥20 reads in the target region was < 95%. We analyzed the variants of each dog separately using UnifiedGenotyper and HaplotypeCaller of GATK and listed all variants detected by either method. We calculated alternative allele frequencies for each variant using SAMtools (ver. 1.6) (5) and created scatter plots of alternative allele frequencies and read counts in all dogs for each variant. We selected variants that clearly showed three or two clusters corresponding to the genotypes upon visual inspection. The genotype of each individual was determined as follows: when the alternative allele frequency was between 0 and 0.15, we assigned 'homozygote' of the reference allele; when the alternative allele frequency was between 0.25 and 0.75, and between 0.85 and 1, we assigned 'heterozygote' and 'homozygote' of the alternative allele, respectively. If the alternative allele frequency was outside these ranges or if a variant position was covered with < 20 sequencing reads, a 'missing genotype' was assigned. All variants were manually inspected in the BAM files using IGV (ver. 2.16.0).

#### Variant annotation and determination of pathogenic variants

Each variant was annotated using SnpEff (ver. 4.3) (6) with the transcripts used in the primer design. Variants with an IMPACT of 'HIGH' in SnpEff were annotated as 'loss of function (LoF),' those with 'MODERATE' were 'nonsynonymous', and those with 'LOW' were 'synonymous'. Because the LoF variants in the 5% terminus of the protein may not lose their function (7), the variants in these regions were not treated as LoF. Because variant annotations vary among isoforms, the annotations for each variant were tallied and the variant annotation with the highest number of annotations was taken as the representative annotation for the variant. If the number of annotations were identical, the annotation assumed to have the weakest impact on the amino acid sequence of the protein was adopted. In other words, the final annotation of variants with the same number of annotations was adopted in the following order: 'synonymous, 'nonsynonymous', and 'LoF'.

In this study, we focused on missense variants on the position that successfully lifted genes from dogs to humans, and adapted only consistent prediction results among three programs to increase specificity: AlphaMissense (8), ESM1b (9), and CADD v1.7 (10). The genomic position of all missense variants was lifted from CanFam3.1 to hg38 using ‘liftover’ function of Picard (v2.26.11). Variants for which position conversion was successfully completed and the amino acids before and after substitution were consistent between dogs and humans were scored for all three programs. When multiple nucleotide variants resulted in multiple amino acid substitutions, the highest score among the individual scores of decomposed amino acid substitutions was adapted. To determine the threshold of the genes analyzed in this study, variants registered in ClinVar (v20240902) (11) with annotation of ‘Pathogenic’, ‘Likely_pathogenic’, ‘Benign’ and ‘Likely_benign’ flagged by the ‘reviewed_by_expert_panel’ or ‘criteria_provided, multiple_submitters, no_conflicts’ in 203 genes were scored using the three programs. The scores with the maximum true positive rate among a false positive rate of 5% or less when predicting the ClinVar annotation were determined for each variant, and the scores were used as the threshold for pathogenicity in this study. Variants that cleared all thresholds for each program were annotated as ‘pathogenic’.

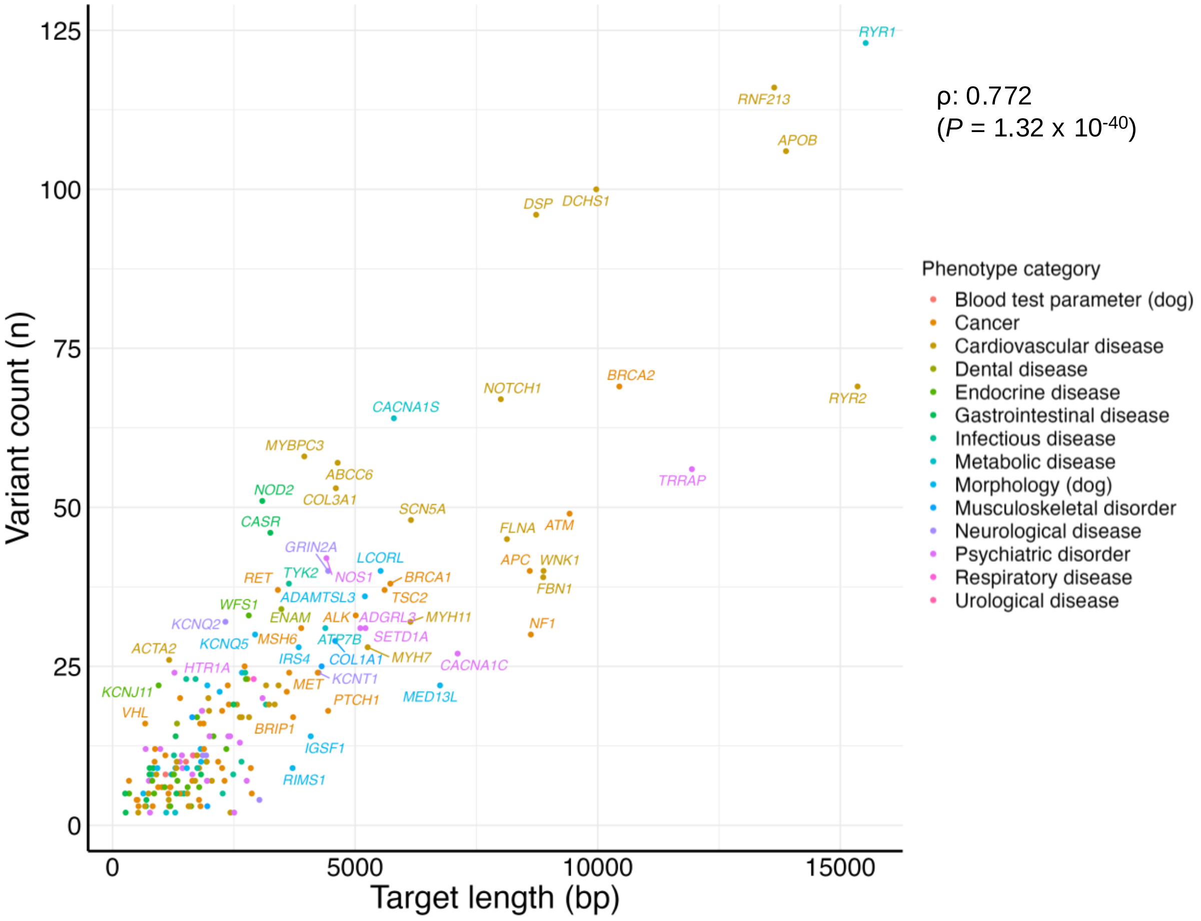


**Figure S1.** The correlation of the length of targeted genes and the number of the variants detected in this study.

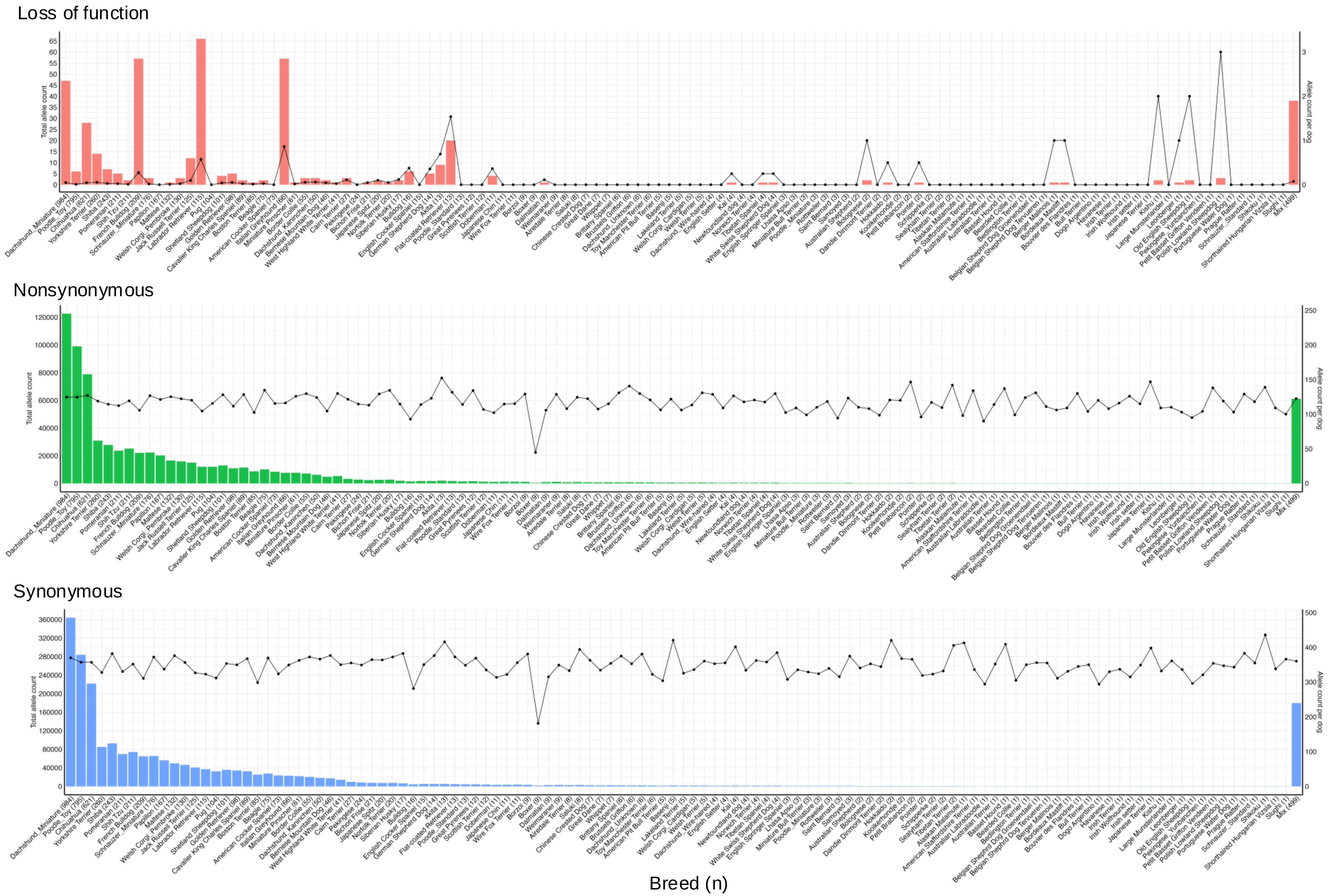


**Figure S2.** The total number of variants detected per breed (bar plot with left y-axis) and the number of alleles per individual for each breed (line plot with right y-axis).

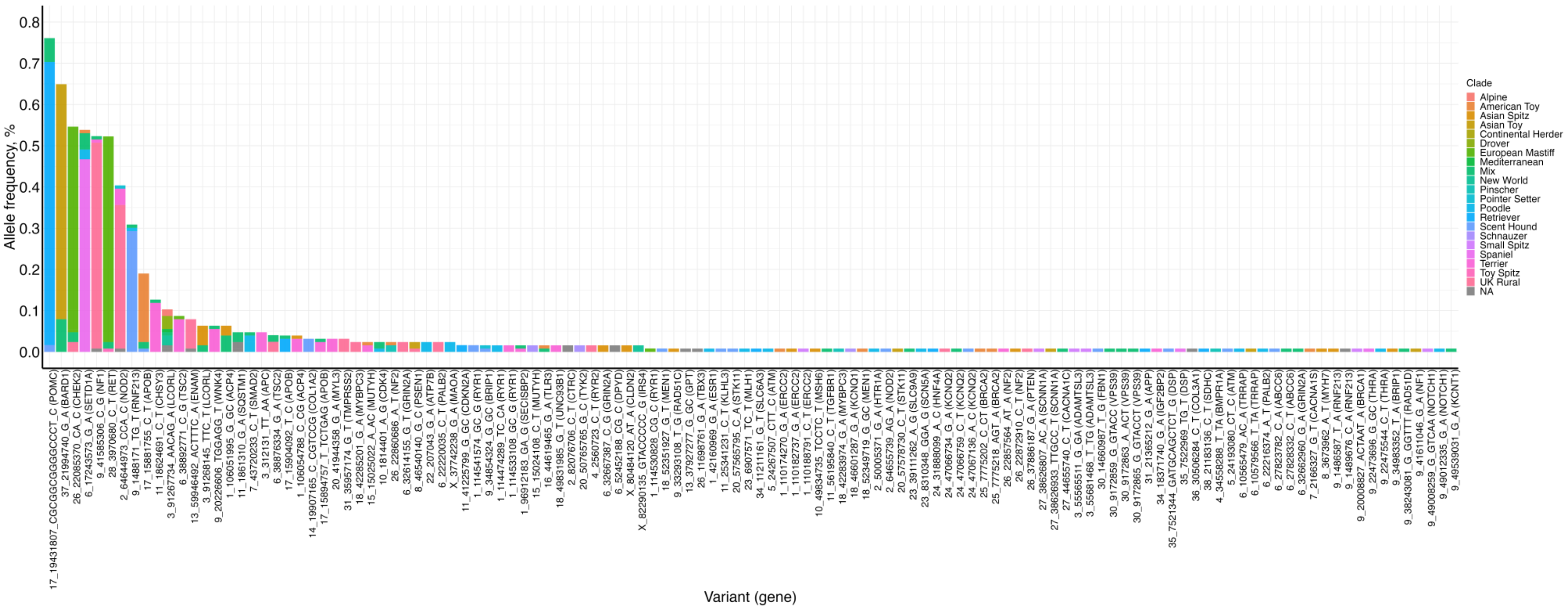


**Figure S3.** The alternative allele frequencies of pathogenic variants in bar plots. The colors of each bar represent the clades to which the breeds of the carrier dogs belong to Parker, et al. (13).

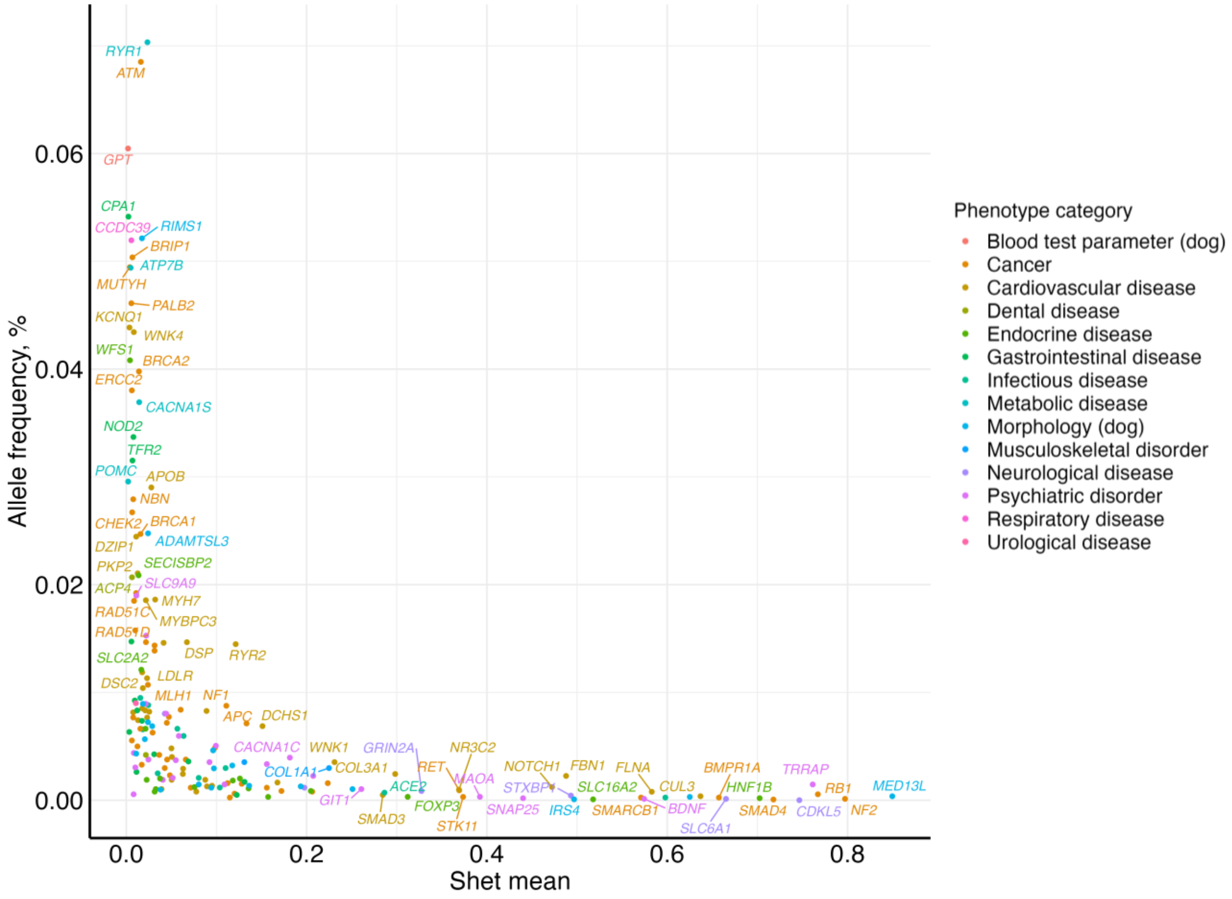


**Figure S4.** Distribution of the alternative allele frequencies in human (gnomAD) and the mean Shet scores of each gene.

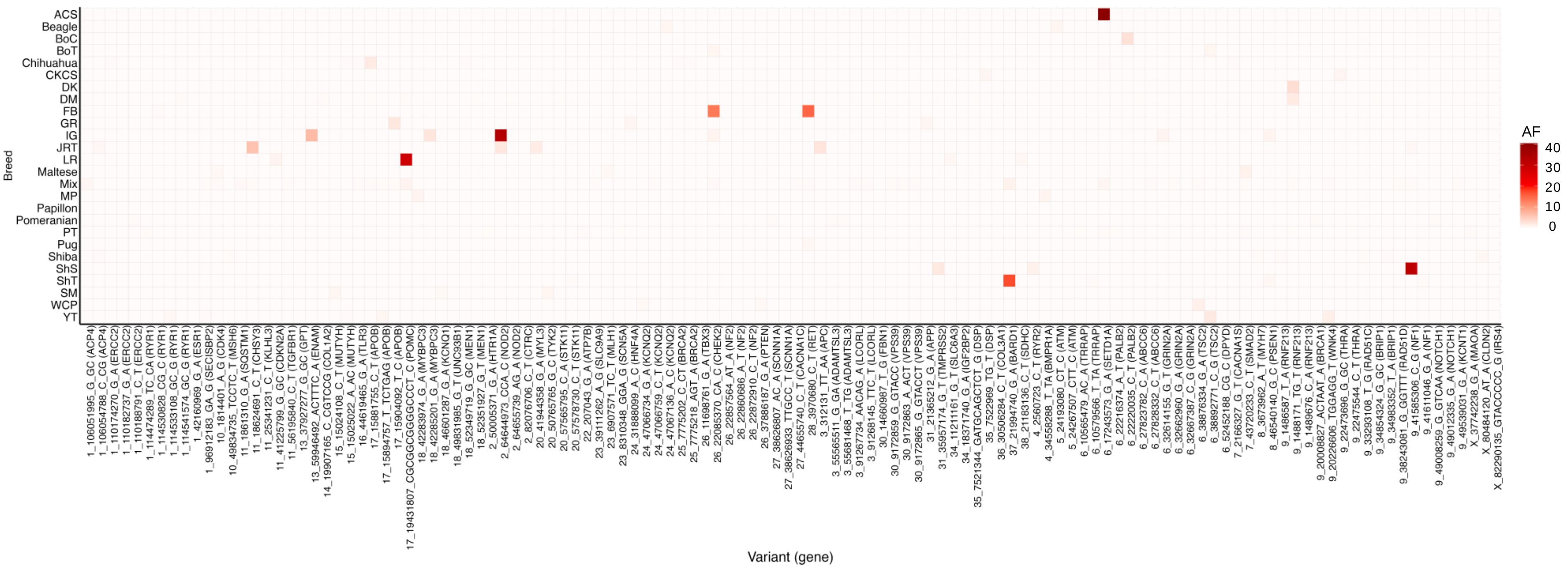


**Figure S5.** The matrix of the allele frequencies of pathogenic variants by breed with over 50 dogs. Abbreviations of breed name are described in Table S2 in SI Appendix. AF, allele frequency.

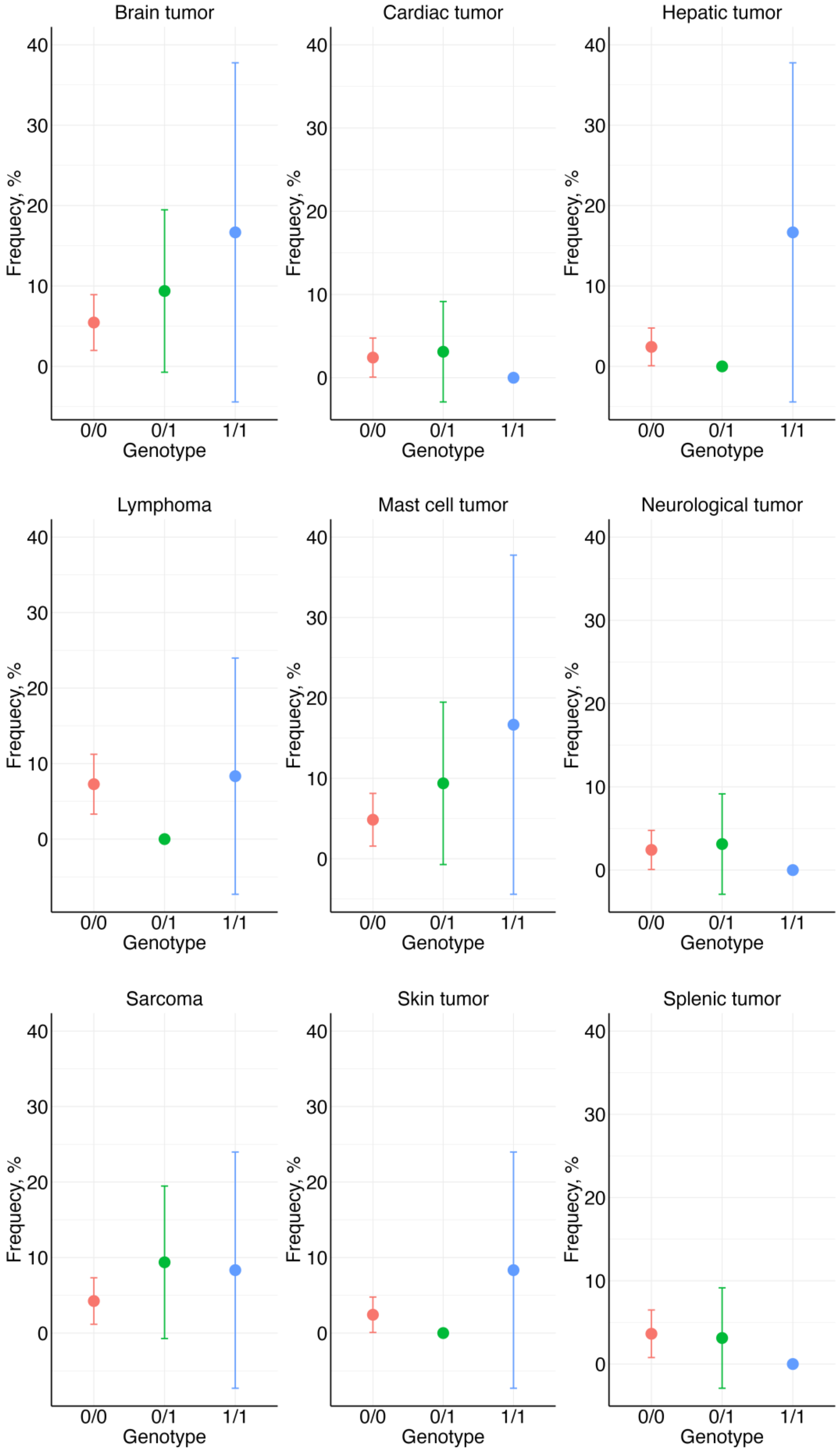


**Figure S6.** Comparison of frequencies of dogs with each cancer among the genotypes of the LoF variant in CHEK2 (p.Leu184fs). Plots and bars represent averages and standard errors of the frequencies. Hepatic cancer showed nominal association (P = 4.00 x 10-2) with the genotype of the variants at the recessive model.

#### Fig. S7.

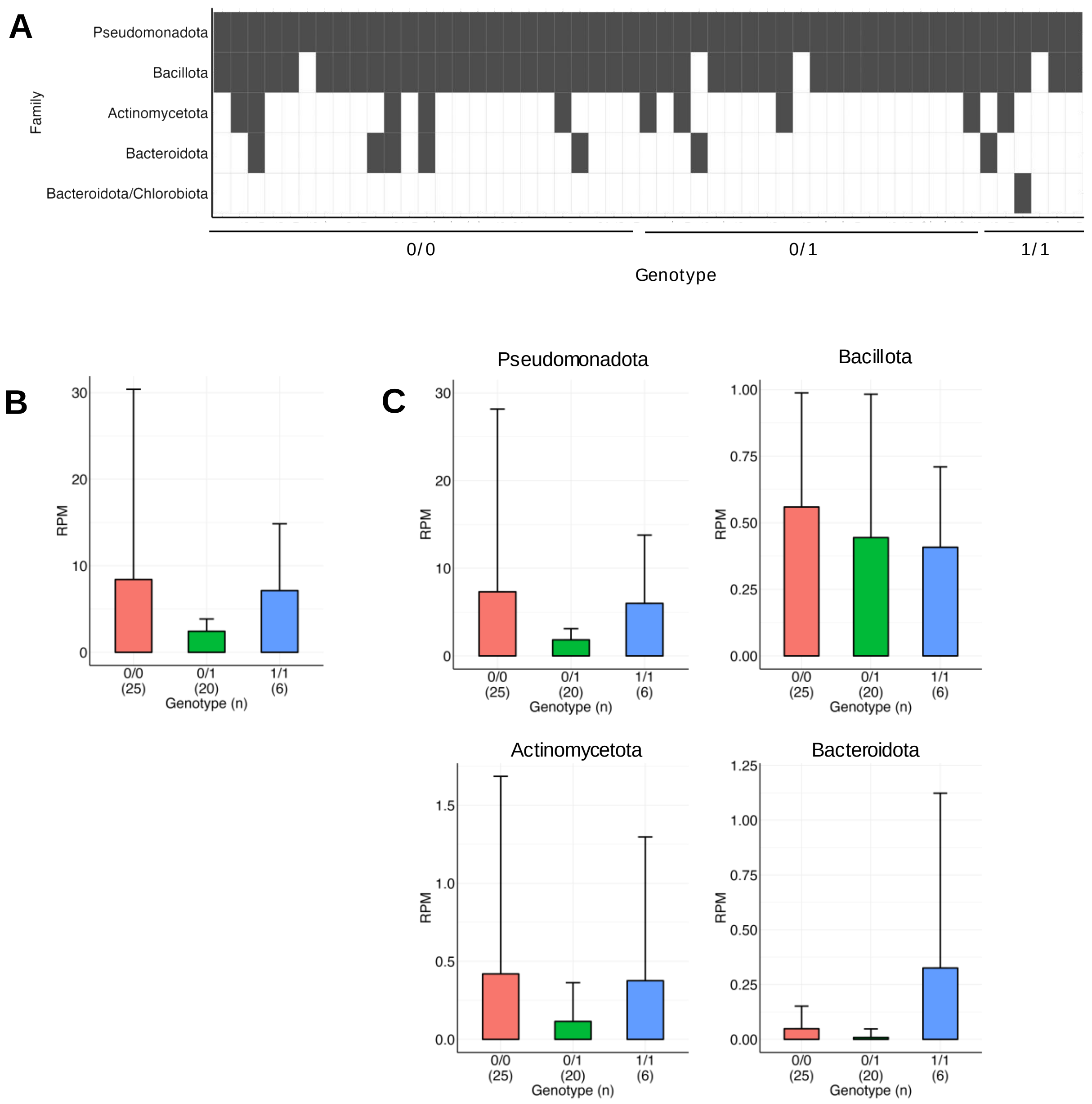


**Figure S7.** The estimated number of bacteria in the peripheral blood for each genotype of the LoF variant in NOD2 using unmapped sequence reads from 51 whole-genome sequencing data. No genotype-dependent alternation in the presence of certain bacteria (A), the total number of bacteria (B) or susceptibility to certain bacteria (C) were observed.

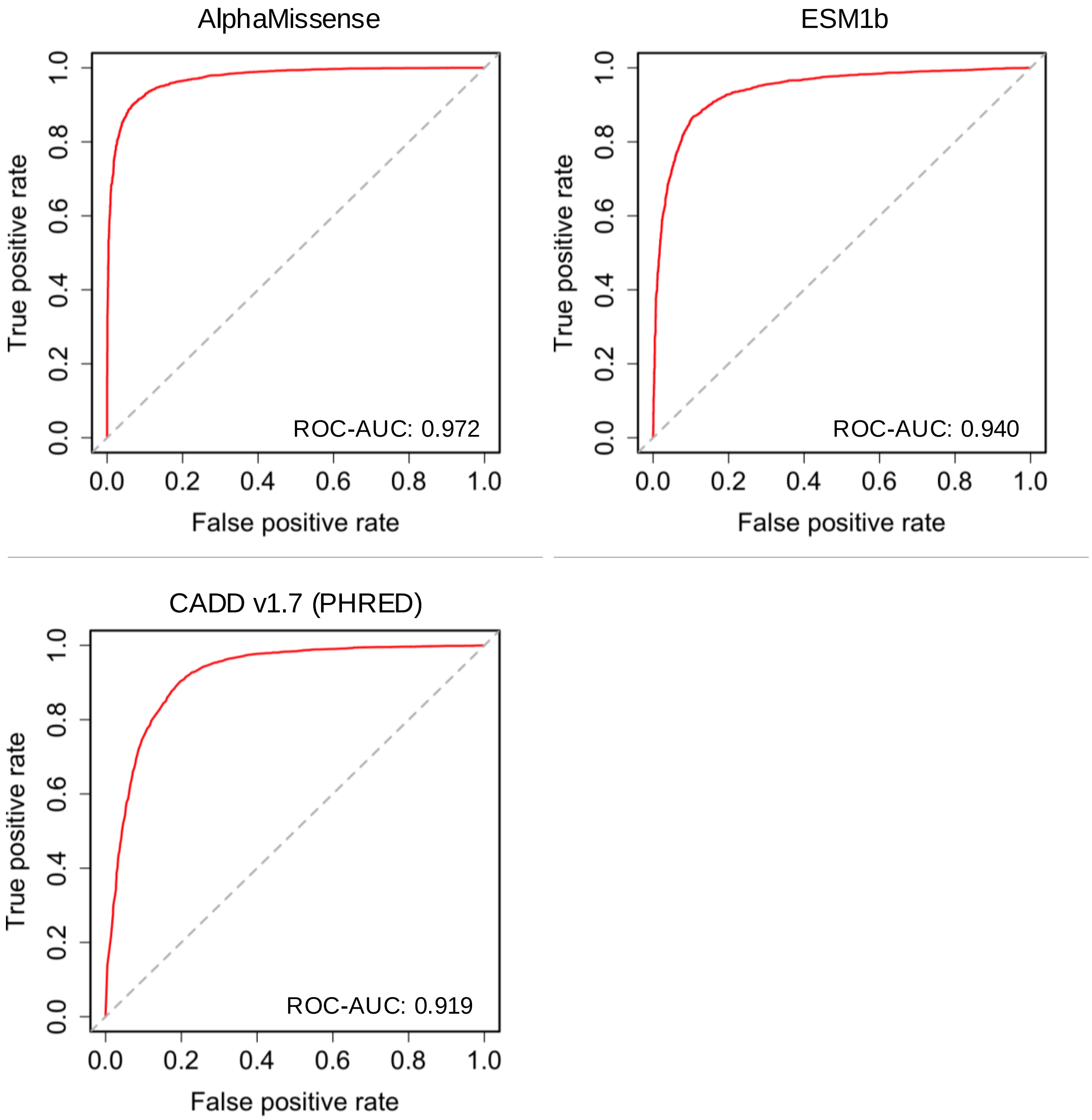


**Figure S8.** Receiver operating characteristic curves showing predictive performance of scores from three in silico programs with ClinVar annotation (pathogenic, likely_pathogenic, benign, and likely_benign) of variants in genes analyzed in this study.
